# Enumerating and Exploring the Space of Clonal Trees

**DOI:** 10.1101/2025.07.18.664776

**Authors:** Bryce Bernstein, Kendra Winhall, Layla Oesper

**Affiliations:** Carleton College, Northfield, Minnesota, USA

**Author notes:** Permission to make digital or hard copies of all or part of this work for personal or classroom use is granted without fee provided that copies are not made or distributed for profit or commercial advantage and that copies bear this notice and the full citation on the first page. Copyrights for components of this work owned by others than the author(s) must be honored. Abstracting with credit is permitted. To copy otherwise, or republish, to post on servers or to redistribute to lists, requires prior specific permission and/or a fee. Request permissions from. Conference’17, Washington, DC, USA. **CCS Concepts** Applied computing →Molecular evolution; Computational genomics; Computational biology. **ACM Reference Format** Bryce Bernstein, Kendra Winhall, and Layla Oesper. 2025. Enumerating and Exploring the Space of Clonal Trees. In. ACM, New York, NY, USA, 10 pages. https://doi.org/10.1145/nnnnnnn.nnnnnnn.

**Keywords:** Tumor Evolution, Clonal Tree, Cancer, Enumeration, Algorithms, Simulated Data

## Abstract

Tumor growth is a complex evolutionary process initiated by an abnormal ancestor cell progressively gaining mutations, which eventually results in uncontrolled cell division. Researchers use a structure called a clonal tree–which depicts ancestral relationships between mutated cell populations–to represent a tumor’s evolutionary history. Numerous methods have been developed to reconstruct a clonal tree from tumor sequencing data. To evaluate the accuracy of such tree inference methods, researchers typically use simulated data where the true evolutionary history is known. However, previous research has not thoroughly analyzed the space of clonal trees under different evolutionary models. Such exploration would help to better understand the underlying structure and characteristics of these spaces of trees and would aid in creation of more appropriate simulated datasets. We analyzed four different categories of clonal trees, each with their own set of assumptions. For each category, we designed and implemented enumeration algorithms that provably generate all such clonal trees with a specified number of mutations. We then used our algorithms to generate several datasets and analyzed the generated trees to discover patterns in the data across different assumptions. We also investigated two tree sampling methods to compare their output with our fully enumerated trees and found that one of the methods does a good job of representing the entire space of trees, while the other does quite poorly. Our findings have important implications for the creation of simulated data commonly used to assess new clonal tree inference methods. Code associated with the project is freely available at: https://bitbucket.org/oesperlab/tree-space/src/main/

## 1 Introduction

Scientists have found that tumors start as a single founder cell which progressively acquires mutations as the cell divides and eventually grows into malignant cancer [23]. Clonal trees, which depict the ancestral relationships between cells with different genotypes, are widely used to represent this evolutionary process. A clonal tree can provide important insight into how the tumor evolved which has the potential for important clinical impacts [12].

Clonal trees are typically inferred from DNA sequencing data, which only captures information about the tumor at one or a few time points, rather than throughout its full history. As such, accurately inferring a clonal tree has proved to be a challenging task. Different clonal tree inference methods utilize different assumptions about how the tumor evolved. For instance, [10, 15, 25] and many other tree inference methods operate under the Infinite Sites Assumption (ISA), which states that once a mutation is gained, it is never lost or gained again [16]. In recent years, tumor evolution models that relax the ISA have become more common, as evidence has suggested that the rigid assumptions of the ISA model are often violated in cancer [18]. The *k*-Dollo model [8] still restricts mutations to only be gained once, but allows for mutations to be deleted up to *k* times. An important restriction on this model is the 1-Dollo model where mutations may be deleted up to one time. The Dollo model (or variants of it) has been utilized by a number of different clonal tree inference methods including [5, 9, 27]. Other models of evolution, such as the Camin-Sokal model [3] which allows for mutations to be gained multiple times (but not deleted) or the finite-sites model which allows for both mutation gains and losses also have been utilized in clonal tree inference algorithms [32, 33]. Many new clonal tree inference algorithms continue to be developed as different kinds of sequencing data become available and different models of evolution or other underlying algorithmic assumptions are utilized to try and solve the challenging problem of accurately inferring tumor evolution [20].

One common task undertaken by virtually all newly developed clonal tree inference algorithms is benchmarking on simulated data where the true underlying clonal tree is known a priori. Recently,[19] undertook a study to evaluate the landscape of different simulation techniques used and found a variety of issues, most notably that the significant majority of papers favor their own custom simulations over tools specifically designed for creating simulations. Another question, not explicitly considered by [19] in their analysis, is how simulations that directly generate clonal trees reflect the entire space of the particular type of clonal trees from which they are sampled. Our work here is motivated by the need to better understand these spaces of clonal trees (an interesting exercise itself) with the aim that this knowledge can be used to improve the creation of simulated data that better reflect the broad spectrum of possible clonal trees under certain evolutionary models.

In this paper we define and analyze four different sub-categories of clonal trees under two different models of tumor evolution: the ISA model and the 1-Dollo model. For each of our four sub-categories we describe an algorithm to enumerate all trees under that model that contain up to a specified number *m* of mutations gained and analyze the resulting algorithms. We also describe two tree sampling approaches for one of our sub-categories of clonal trees. We use our enumeration algorithms to create 4 datasets of enumerated clonal trees and analyze the features of these datasets using 7 different measures that capture both topology and label-based aspects of the clonal trees. We also create two datasets using our sampling procedures and compare the results to the fully enumerated dataset. In this case, we find that one of the sampling procedures, which has been commonly used in the creation of simulated data, does not accurately reflect the entire space of clonal trees. This finding has important implications for how clonal tree simulations may be done in the future.

## 2 Methods

The evolutionary history of a tumor is often represented using a rooted, node-labeled tree. Each vertex in the tree represents a *clone*, or a set of cells that share the same somatic mutations, that existed at some point during the tumor’s development. Edges represent ancestral relationships between those clones. Node labels indicate mutations that appeared (or were deleted) in the corresponding clone. All mutations, unless they are explicitly deleted, are inherited by all descendant clones. Broadly, we will refer to the set of all such trees as *clonal trees*.

Different models of evolution, or assumptions about how a clonal tree evolves over time, are often applied to obtain sub-categories of these clonal trees. In sections 2.1-2.4 we use two models of evolution to define four such sub-categories of clonal trees: (1) single-label ISA trees; (2) multi-label ISA trees; (3) restricted 1-Dollo trees; and (4) 1-Dollo trees. Figure 1 contains an example clonal tree for each sub-category. Furthermore, we describe an algorithm for enumerating the space of all trees in each sub-category containing a specified number *m* of mutations gained. We also provide analysis on the efficiency of these algorithms and the size of the space of trees being enumerated. Then in section 2.5 we describe two methods for sampling from the space of single-label ISA trees.

**Figure 1:**
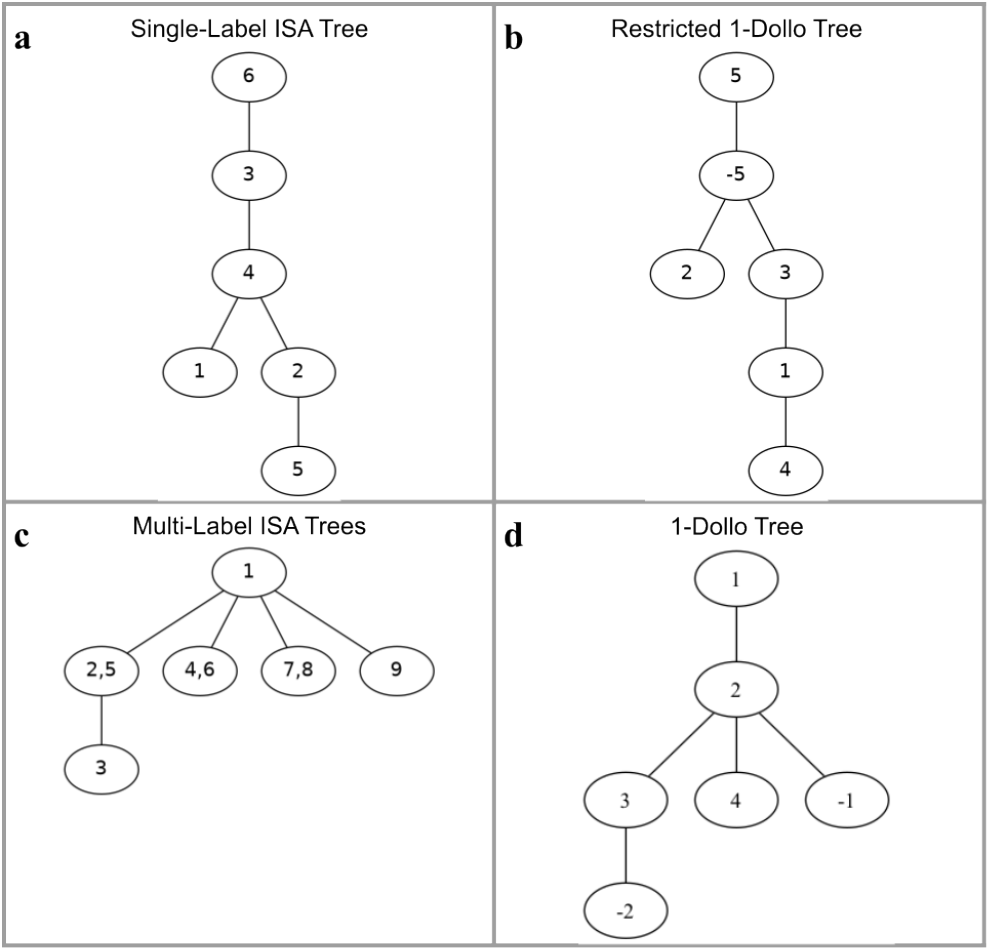
Examples of four sub-categories of clonal trees. (a) A single-label ISA tree containing 6 mutations. (b) A restricted 1-Dollo tree where 5 mutations are gained and the mutation 5 is lost once (as indicated by the negative sign). (c) A multi-label ISA tree containing 9 mutations across 6 clones. (d) A 1-Dollo tree where 4 mutations are gained and mutations 1 and 2 are each lost once (as indicated by the negative signs).

### 2.1 Single-label ISA Trees

The Infinite Sites Assumption (ISA) [16] states that all mutations gained are never lost, and that no mutation is gained more than once during the development of the tumor. This assumption has historically been a common assumption among the tumor evolution research community. An additional assumption that all acquired mutations can be explicitly ordered (so no more than 1 new mutation appears in any clone) has also been commonly used [9, 15]. One way this has been applied is by clustering mutations into clones prior to inferring clonal trees [2]. We therefore define a *single-label ISA tree* to be a clonal tree that adheres to both of these assumptions (see Figure 1a for an example). We note that in the literature these trees are often referred to as *mutation trees* [1, 15], but chose to use the name single-label ISA trees here to highlight the underlying assumptions defining this sub-category of clonal trees. Furthermore,we define the set ℐ_*m*_ to be the set of all single-label ISA trees where *m* unique mutations have been gained.

#### 2.1.1 Enumerating Single-label ISA Trees

We describe here an algorithm to enumerate the set of all single-label ISA trees with *m* mutations, or ℐ_*m*_. This algorithm utilizes something called a *Prüfer sequence*–a sequence of length *m*−2 containing values (with repetition allowed) from the set {1, …, *m*} [24]. It is well known that there is a bijective relationship between Prüfer sequences of length *m*−2 and *unrooted* trees on *m* labeled vertices. Algorithm 1 describes how to do this conversion. In short, edges are iteratively added to the unrooted tree such that each edge added has one end connected to the next element in the Prufer sequence and the other end is connected to the smallest remaining node value that hasn’t been connected to in a previous iteration, and that does not appear in the remaining portion of the prufer sequence. Finally, the two remaining nodes that haven’t been connected to previously with a node from the prufer sequence are linked together. See Figure 2 for an example execution of Algorithm 1.

**Figure 2:**
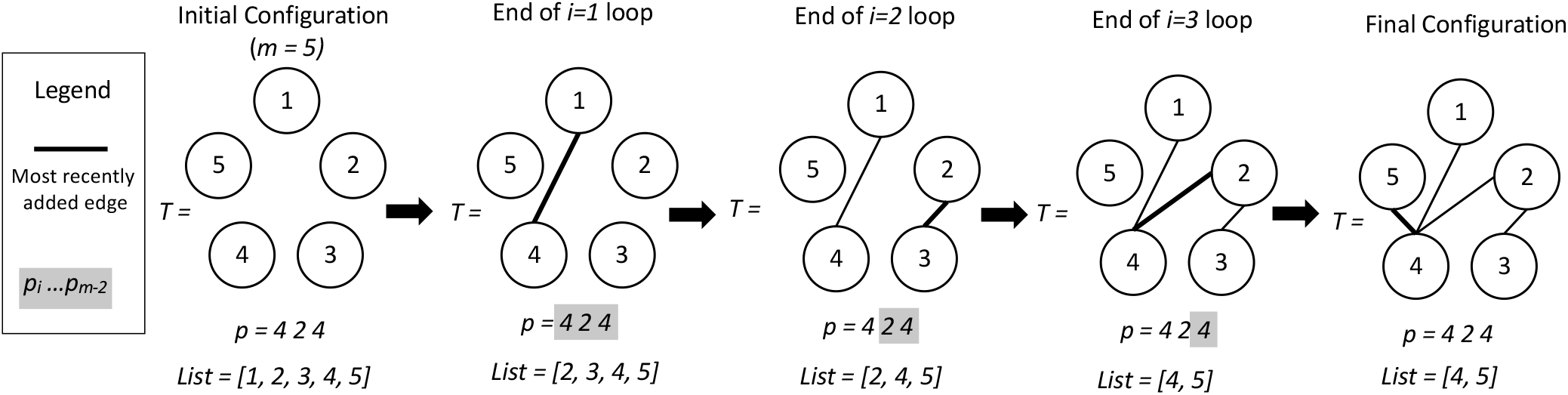
An example demonstrating the execution of Algorithm 1 to construct an unrooted tree with *m* = 5 labeled nodes from the Prüfer sequence p = 424.

We can therefore enumerate ℐ_*m*_ with the following steps: (1) Enumerate all Prüfer sequences of length *m*−2; (2) Use Algorithm 1 to convert each Prüfer sequence into the corresponding unrooted tree *T* with *m* labeled nodes; and (3) Create *m* rooted trees from each unrooted tree *T* by choosing each vertex of *T* as the root of a separate tree.

##### Algorithm 1

Turn Prüfer Sequence into Unrooted Tree with *m*Labeled Nodes

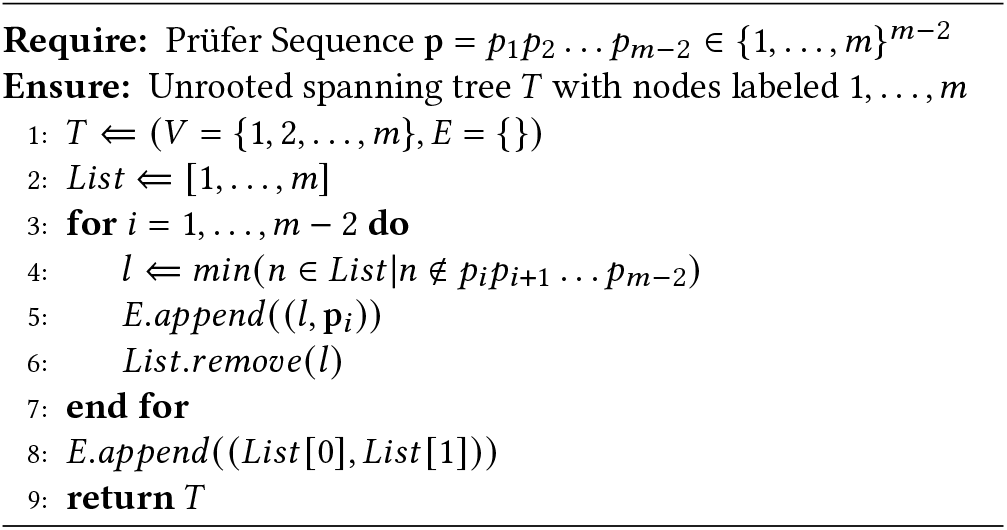

#### 2.1.2 Analysis of Single-label ISA Algorithm

Algorithm 1 can be implemented as efficiently as *O* (*m*) [30]. Rooting the output tree at all *m* nodes takes *O*(*m*), thus making the complexity to turn a single Prüfer sequence into a set of *m* rooted ISA trees *O*(*m*). According to Cayley’s formula, there are *m*^*m*−2^ unrooted trees on *m* vertices [4]. Thus, there are *m*^*m*−2^ Prüfer sequences of length *m*−2 because of the previously mentioned bijective relationship. This means that |ℐ_*m*_ | = *m*^*m*−1^ since each unrooted tree has *m* rooted trees and ℐ_*m*_ can therefore be fully enumerated in *O*(*m*^*m*−1^) time. To understand the scale of this growth, note that ℐ_13_, the set of single-label ISA trees on 13 mutation, contains more than 23 trillion trees. This is more than the number of physical trees in the real world (an estimated 3.04 trillion) [6].

### 2.2 Multi-label ISA Trees

We define a *multi-label ISA tree* as a rooted, node-labeled tree that adheres to the ISA, but drops the assumption that all mutations can be explicitly ordered. As a result, multiple mutations may label a single node–indicating that the exact order these mutations were gained cannot be inferred. Note that the set of single-label ISA trees is a subset of the set of multi-label ISA trees. See Figure 1c for an multi-label ISA tree. We note in the literature these trees are often called *clonal trees*, but chose to use the name multi-label ISA tree here to make it clear exactly how these relate to single-label ISA trees defined in section 2.1.

#### 2.2.1 Enumerating Multi-label ISA Trees

Let 𝒞_*m*_ be the set of all multi-label ISA trees with 𝒞 *m* mutations. Trees in this set can have up to *m* nodes (in this case, they exactly correspond to the set of single-label ISA trees). Therefore, we will find it useful to further sub-divide this set. We define 𝒞 _*m,n*_ be the set of all multi-label ISA trees with *m* mutations and *n* nodes, where 0 *< n*≤*m*. Lastly, we define *L*_*m,n*_ to be the set of all partitions of the mutations 1, 2, …, *m* into *n* distinct sets. For example, 1, 2|3|4|,5,6 and 1, 4, 6 2 3 5 are both elements of *L*_6,4_. Algorithms exist to enumerate all elements in *L*_*m,n*_ [17].

Algorithm 2 describes how to enumerate all trees in _*m,n*_ for fixed variables 𝒞*m* and *n*. In short, start with all single-label ISA trees with *n* mutations, or ℐ_*n*_. Note that these trees have exactly *n* nodes because they are single-label. We then apply each mutation partition in *L*_*m,n*_ to the corresponding nodes in each single-label ISA tree to construct a tree with *n* nodes and *m* mutation labels.

##### Algorithm 2

Enumerate the Set of All Multi-label ISA Trees with *m* Mutations and *n* Nodes

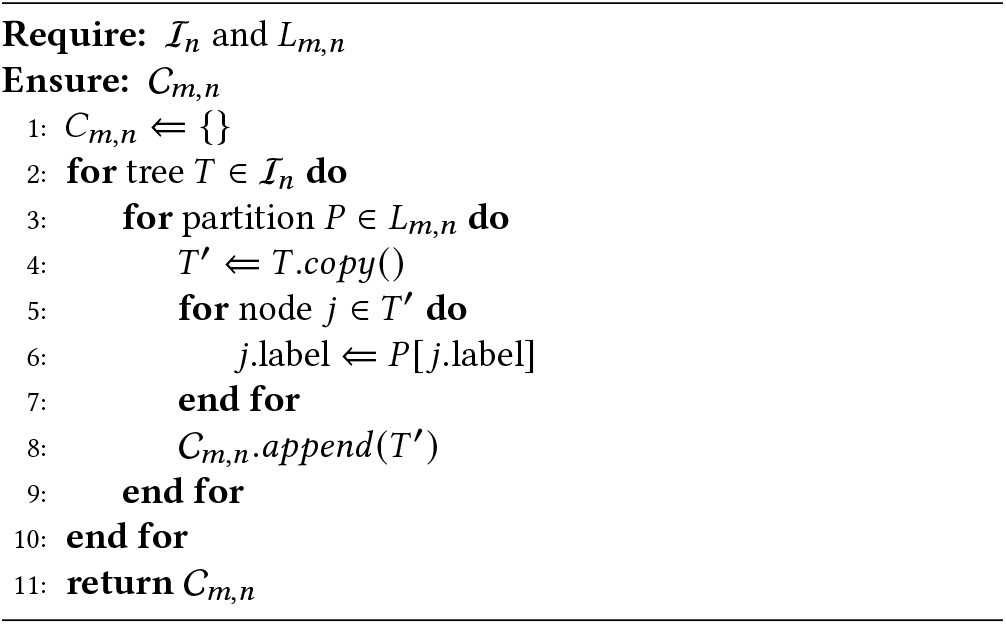

Finally, we can use Algorithm 2 to enumerate ℐ_*m*_ by concatenating its output for *n* = 1, …, *m* into a single set of trees. That is, 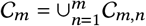

#### 2.2.2 Analysis of Multi-label ISA Algorithm

As discussed in the previous section, |ℐ_*n*_| = *n*^*n*−1^, so the outer most loop of Algorithm 2 executes *n*^*n*−1^ times. The number of partitions of *m* distinct objects into *n* indistinguishable sets, or |*L*_*m,n*_|, is given by *S* (*m, n*), the Stirling number of the second kind [14]. Thus, the middle loop of Algorithm 2 executes *S* (*m, n*)times, giving us | 𝒞_*m,n*_ | = *S* (*m, n*)∗*n*^*n*−1^. Finally, the innermost loop executes *n* times, so Algorithm 2’s runtime is *O* (*n*^*n*^ ∗ *S* (*m, n*)).

Enumerating all trees in 𝒞_*m*_ is done by iteratively applying Algorithm 2 with *n* = 1, …, *m* and concatenating together the results. Thirs means that that the runtime to generate 𝒞 _*m*_ can be written as 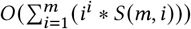

### 2.3 Restricted 1-Dollo Trees

Recent research has shown that the ISA is often violated [18] in tumor evolution, so researchers have begun exploring other models. For example, the *k*-Dollo model allows for each mutation to be deleted at most *k* times in one tree [11]. The number of trees for even the 1-Dollo model rapidly grows restrictively large, so we start by describing a particular subset of these trees. We define a *Restricted 1-Dollo Tree* to be a tree where: (i) Each mutation can evolve at most once; (ii) A *single* mutation is deleted *exactly* once. This adds some flexibility to the ISA model while maintaining many of its rules. See Figure 1b for an example of a restricted 1-Dollo tree.

#### 2.3.1 Enumerating Restricted 1-Dollo Trees

Let 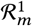 be the set of restricted 1-Dollo trees with *m* mutations gained (and exactly one mutation lost a single time). To generate 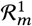 we start from all single-label ISA trees with *m* mutations, or ℐ _*m*_. For each *T* ∈ ℐ _*m*_ and each vertex *v* in *T* we create a set of trees that have the deleted mutation as a child of *v*. Selecting how to insert this child requires enumerating all possible subsets of the current children of *v* and making the selected subset become children of the inserted node. The unselected nodes remain as siblings of the inserted node. Finally, the identity of the deleted node can be any of the mutations on the path from the root to node *v*. Algorithm 3 describes this complete procedure and Figure 3 shows an example of all possible mutation deletions insertions for a given single-label ISA tree. Note that in Algorithm 3, operations occurring inside the inner most for loop that involve *C* or *v*, refer to the corresponding nodes in *T* ^′^.

**Figure 3:**
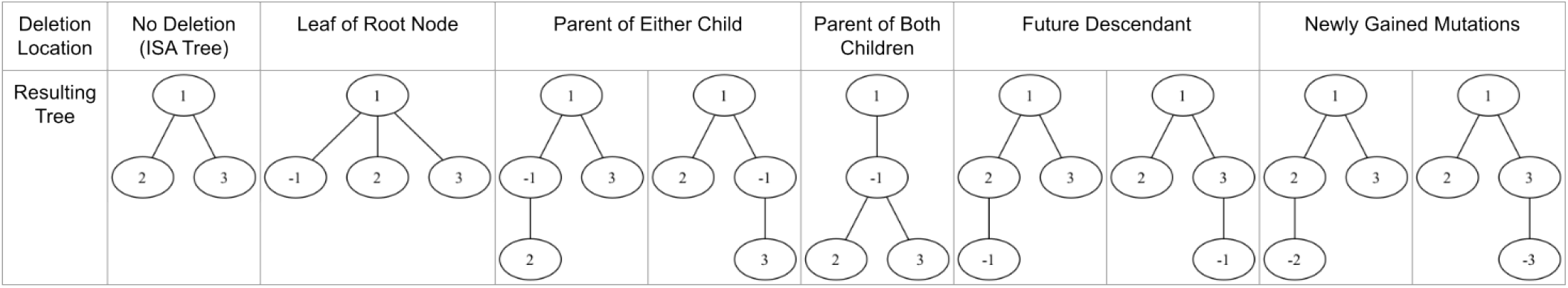
All the possible insertions of a deletion to the single-label ISA tree on the far left, as performed by Algorithm 3. Each panel represents a unique tree that is created, grouping trees created using similar patterns of insertions.

#### 2.3.2 Size of the tree space

Each single-label ISA tree *T* in ℐ_*m*_ generates a number of restricted 1-Dollo trees for each node *v* in *T*. The number of trees generated is determined by the characteristics of the node *v* being considered. When d(*v*) represents the depth of *v* and *c* (*v*) represents the number of children *v* has, the process creates d(*v*)∗ 2^*c* (*v*)^ restricted 1-Dollo trees. We get d (*v*) because, in a single-label environment, depth is equivalent to number of mutations acquired and any acquired mutation can be deleted. Moreover, a tree must also be created for each combination of *v*’s children (see Figure 3) to generate all trees, so there are 2^*c* (*v*)^ trees seen here. Thus, we reach our conclusion that the number of created trees by a node *v* is d *v* 2^*c* (*v*)^. So, for any tree *T* = (*V, E*) ∈ℐ_*m*_,*T* will generate the following number of restricted 1-Dollo trees: 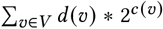 .This is done for all | ℐ_*m*_| = *m*^*m*−1^ single-label ISA trees.

#### 2.3.3 Analysis of Restricted 1-Dollo Algorithm

The process to insert a single deletion into a particular position in an existing single-label ISA tree can be accomplished in linear time, but the process of turning one single-label ISA tree into all consequent restricted 1-Dollo trees cannot. The space of restricted 1-Dollo trees grows exponentially. Interestingly, a single-label ISA tree with 4 or more mutations gained can generate at most 2+2^*m*−1^ +3∗(*m* − 2)restricted 1-Dollo trees as we prove below.

##### Algorithm 3

Enumerate all restricted 1-Dollo trees with *m* mutation gains

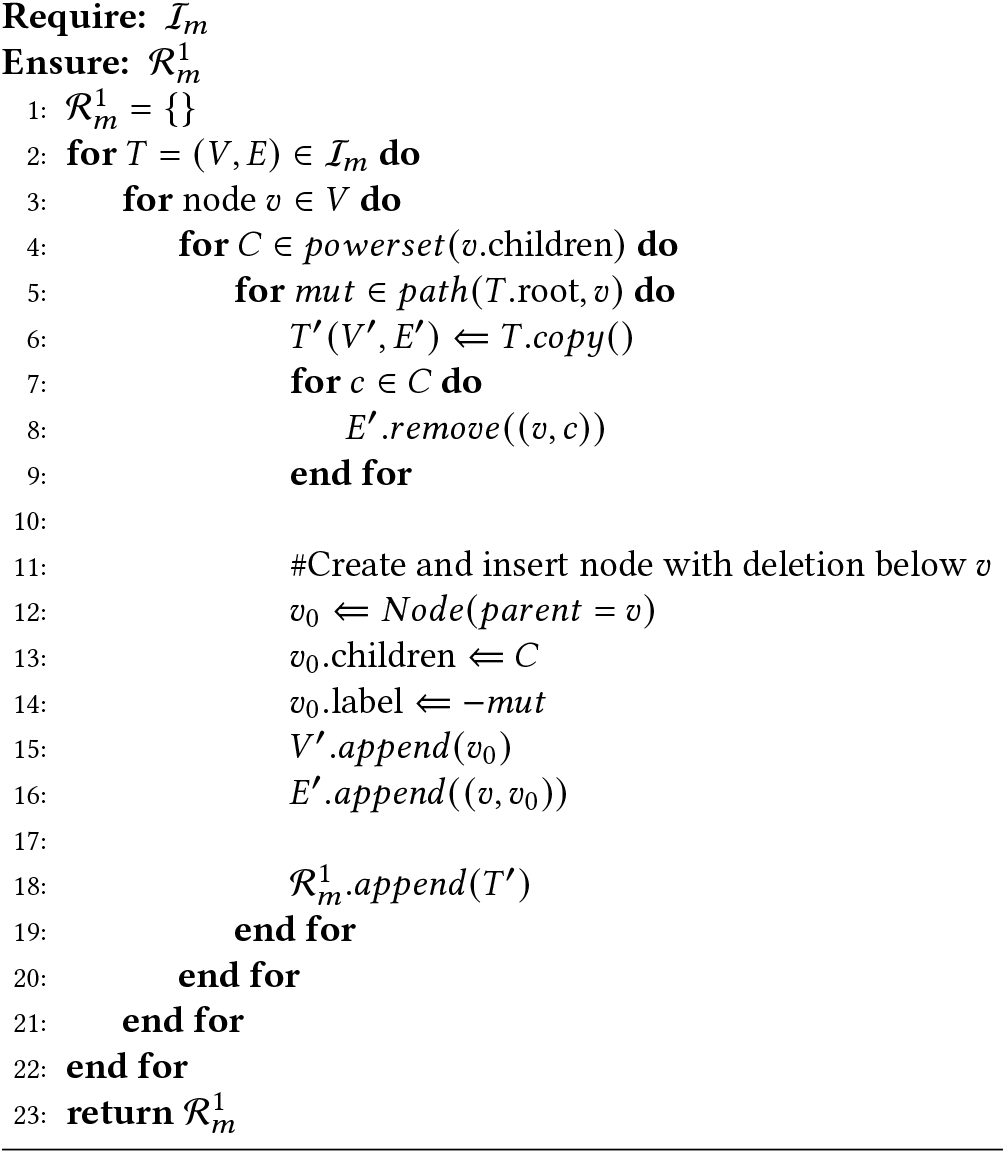

##### PROPOSITION 2.1.

*The maximum number of restricted 1-Dollo Trees generated from an ISA Template tree is* 2 + 2^*n*−1^ + 3(*n* − 2) *for n ≥* 4.

PROOF. We build on the fact that any individual node creates d (*v*) ∗2^*c* (*v*)^ to show this. Notice that this formula consists of a linear and an exponential term. We know that exponential terms grow faster, so, intuitively, our interest will be in maximizing this term. This leads us to the tree that has a root node with all other nodes as children of that root. From the root node, we can conclude that it generates 1 ∗ 2^*n*−1^ trees. Each of the *n* − 1 leaves then generates 2 trees. This tree, therefore, generates 2^*n*−1^ + 2(*n* − 1) trees. We observe that this formula is greater than 2 + 2^*n*−1^ + 3(*n* − 2) for all *n<* 4.

Increasing the number of mutations gained does not affect our emphasis on increasing the exponential term in our equation, so we must continue maximizing this term. However, we can see that a structure of a root with 1 child that has all other nodes as its child will double the depth of the exponential node from 1 to 2. This results in 1 ∗ 2^1^ being the number of trees generated by the root node, 2 ∗ 2^*n*−2^ trees being generated by its child, and 3 ∗ 2^0^ nodes for all leaves. Notice that 2 ∗ 2^*n*−2^ = 2^*n*−1^, meaning we cancel out the effect of increasing the depth of the exponential node by taking advantage of the doubling depth. This results in the tree generating 2+2^*n*−1^ ∗3(*n*− 2)1-Dollo Trees, our claimed maximum. We will show that further increases in the exponential depth will not increase the number of trees generated.

We can see that further increasing the depth would result in depth going from 2 to 3, but this is not a doubling. Because the depth has not doubled, the change cannot offset of taking one child away from the exponential node. Thus, this change does not increase the number of trees generated.

We can therefore conclude that the maximum number of restricted 1-Dollo trees an ISA tree can create is 2 + 2^*n*−1^ + 3(*n* −2) □

Therefore, generating all the restricted 1-Dollo trees that originate from a single ISA tree is *O*(*m* ∗ 2^*m*−1^). With *m*^*m*−1^ such ISA trees, the overall process has runtime by *O*(2^*m*−1^*m*^*m*^). Interestingly, we note that the total number of restricted 1-Dollo trees with *m* mutations gained matches a sequence found in Sloane [28], which is named as the normalized total height of all nodes in all rooted trees with *m* labeled nodes.

### 2.4 1-Dollo Trees

While the restricted 1-Dollo approach is valuable, we can expand it further to allow multiple deletions in one tree. In the *1-Dollo model*, every mutation in a tree can be deleted up to one time. This model accounts for mutations that occur in a tumor but are subsequently lost–a process that occurs frequently in tumor evolution [18]. See Figure 1d for an example of a 1-Dollo tree with 4 mutations gained and 2 mutations lost.

#### 2.4.1 Enumerating 1-Dollo Trees

First we define 𝒟_*m*_ to be the set of all 1-Dollo trees with *m* mutations gained. Our goal is to enumerate 𝒟_*m*_, but to do so we will need to consider some notation for a particular subset of this set. Let *M* be some subset of mutations in {1, 2, …, *m*}. We define 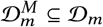 to be the subset of 1-Dollo trees where any deletion appearing is for a mutation from the set *M*.

We first define a helper algorithm that takes in as input some subset of 1-Dollo trees 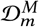 and some mutation *mut* ∈ {1, 2, …, *m*} but where *mut* ∉ *M*. This algorithm works very similarly to the restricted 1-Dollo generation algorithm, or Algorithm 3, by using 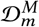 as a set of template trees and inserting a deletion to mutation *mut* in all possible places in all trees in that set. This algorithm then returns 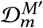 where *M*^′^ = *M* ∪ {*mut*} and is shown below as Algorithm 4.

##### Algorithm 4

Generate 1-Dollo trees by deleting mutation *mut* from trees in 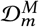

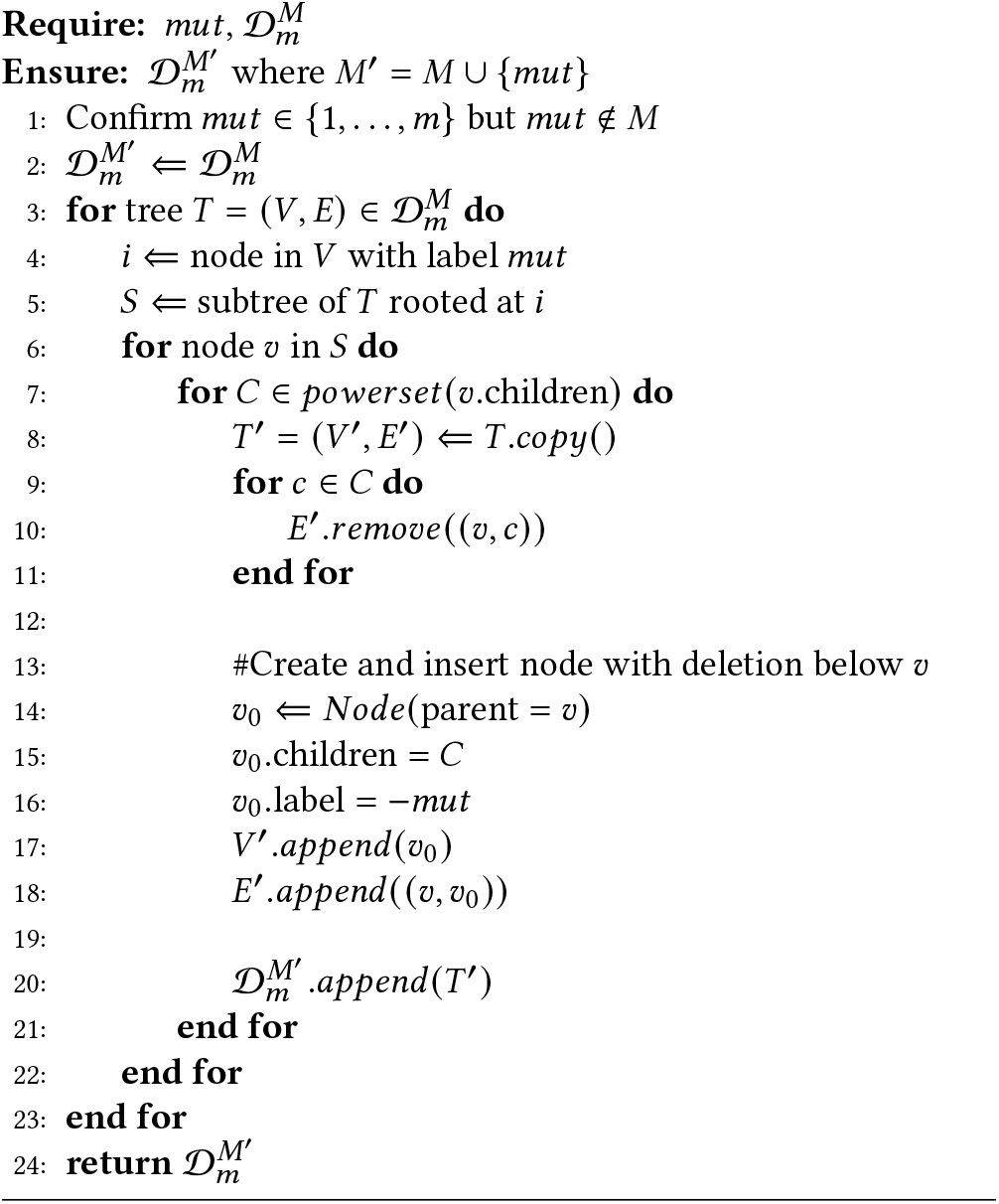

We can now describe how to use our helper Algorithm 4 to enumerate 𝒟_*m*_. We will start with ℐ_*m*_ which happens to be the same thing as 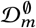 (the set of 1-Dollo trees where no mutations have yet been deleted) and will iterative add mutations 1, 2 …, *m* in that order to construct 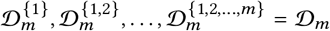. This is shown below as Algorithm 5.

#### 2.4.2 Analysis of 1-Dollo tree Algorithm

While we don’t have a formal theoretical analysis of its runtime, the size of the space of 1-Dollo trees appears to grow quite quickly. We found that there is 1 1-Dollo tree with 1 mutation gained, 18 1-Dollo trees with 2 mutations gained, 645 1-Dollo trees with 3 mutations gained, 42080 1-Dollo trees with 4 mutations gained, 4404375 1-Dollo trees with 5 mutations gained, and 673777200 1-Dollo trees with 6 mutations gained.

##### Algorithm 5

Generate 𝒟_*m*_, all 1-Dollo trees with *m* mutations gained

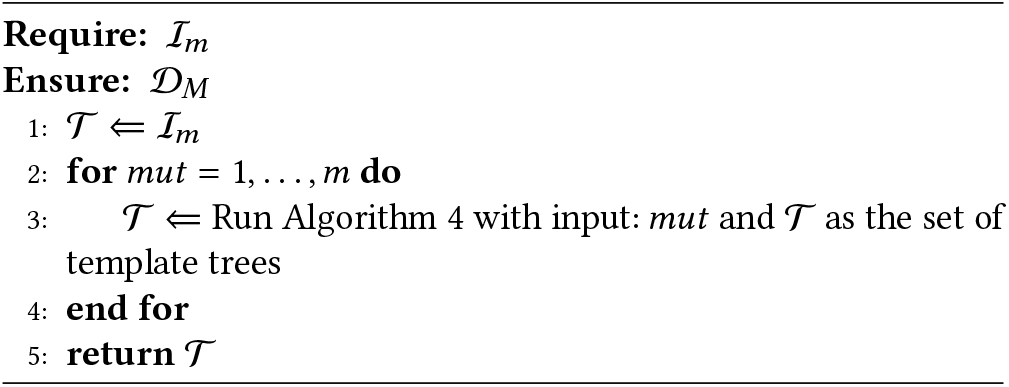

### 2.5 Sampling Clonal Trees

Much of the previous subsections have been dedicated to describing methods that are able to enumerate all clonal trees in certain sub-categories for a specified number *m* of mutations gained. However, as the number *m* of mutations gained increases, these methods quickly become intractable. We therefore pivot here to briefly consider approaches for sampling (ideally uniformly) from these spaces of clonal trees rather than trying to enumerate all trees. For the remainder of this section, we restrict our consideration to the simplest clonal tree sub-category: single-label ISA trees (although our approaches could be extended to the multi-label ISA trees as well). We describe below two different methods for sampling from the space of single-label ISA trees.

#### 2.5.1 A Psuedorandom Method

An simple approach for sampling single-label ISA trees is to iteratively build a tree by adding new mutations by randomly selecting an existing node to be the parent for the added node. A number of papers have used this method for constructing simulated clonal tree data [5, 10, 13, 15, 21, 29]. While on the surface this seems like a reasonable way to sample from the space of single-label ISA trees, it turns out it does not in fact sample uniformly from the space of single-label ISA trees with *m* mutations gained. We show this empirically in section 3.3. We are also able to argue this theoretically by calculating the probability of generating a linear tree with mutation 1 at the root and mutation *m* as the single leaf using this approach and showing that this is not equal to 1 divided by the number of single-label ISA trees with *m* mutations gained. We call this approach for sampling clonal trees the *pseudorandom* approach.

#### 2.5.2 Wilson’s Algorithm

Wilson’s algorithm [31] is an approach that is known to sample uniformly from the space of random rooted spanning trees, which means we can use it to sample uniformly from the space of single-label ISA trees by treating vertex IDs as mutations. In short, this algorithm works by performing loop erased random walks from a node not yet in the tree until a node that is in the tree is encountered. All nodes on the path from the start to the end in the walk are then added to the tree in that order. We note here that while we only consider single-label ISA trees sampled using Wilson’s algorithm, it can be modified to also sample uniformly from the space of multi-label ISA trees. While this approach is guaranteed to sample uniformly, we have only been able to find one paper that uses this approach for the creation of simulated clonal trees [26].

## 3 Results

In section 3.1 we first describe 4 datasets we created using the Algorithms described in sections 2.1-2.4. Then in section 3.2 we provide a thorough analysis of the datasets we created, including expected and unexpected findings. Finally, in section 3.3 we describe how we used the sampling algorithms described in 2.5 to create two new datasets and compare them to the correspond fully enumerated dataset - finding that only one of our sampling algorithms accurately represents the entire space.

### 3.1 Enumerated Dataset Creation

We used our enumeration algorithms described in the previous section to construct 4 datasets. Each dataset covers the space of specific sub-category of clonal trees for a specified range of number of mutations gained. This includes datasets for the following: (1) single-label ISA trees, (2) multi-label ISA trees, (3) restricted 1-Dollo trees, and (4) 1-Dollo trees (see Table 1 for a summary). In order to highlight differences between these different classes, we did not include all single-label ISA trees as part of the last three categories in our dataset construction. That means in our dataset of multi-label ISA trees only contains the trees in this set that are not also single-label ISA trees (the same is true for the restricted 1-Dollo and 1-Dollo datasets). The upper-bound on the number of mutations gained in our simulated datasets was dictated by the running time to produce the dataset. For example, for the single-label ISA tree dataset we were able to generate all such trees that contained up to 11 unique mutations gained, but for the 1-Dollo dataset we could only create all trees that contained up to 6 unique mutation gains. In most cases we constructed all trees with number of mutations gained from 1 to our upper-bound. The one exception is the multi-label ISA trees where we stared with 2 mutations since that was the number of mutations required for the tree to be a multi-label ISA tree, but not a single-label ISA tree. In total we created over 108 billion unique clonal trees.

**Table 1:** A summary of the datasets created using our enumeration algorithms for different sub-categories of clonal trees.

| Type of Clonal Tree | # Mutations Gained | Total # Trees Generated |
| --- | --- | --- |
| Single-label ISA | 1-11 | ~27 billion |
| Multi-label ISA | 2-10 | ~4.5 billion |
| Restricted 1-Dollo | 1-10 | ~76 billion |
| 1-Dollo | 1-6 | ~700 million |

In the following subsection, we use these datasets to explore features and uncover patterns of these spaces of clonal trees.

### 3.2 Analysis of Enumerated Datasets

We analyzed our enumerated datasets using 7 different topological and label-based measurements.

#### 3.2.1 Tree Measurements

Our topological measurements include tree height; minimum, maximum, and average branching factor; and number of leaves. Specifically, tree height measures the number of edges between the root and the furthest node. Minimum branching factor measures the smallest number of children that a non-leaf node has, maximum branching factor measures the largest, and average branching factor averages the branching factor of all non-leaf nodes in the tree. Finally, the number of leaves for a tree counts the number of nodes without children.

We also considered two label-based measurements. In a clonal tree, each node is labeled with the name of mutations first appearing in the corresponding clone. So, unless a mutation is deleted, each clone inherits all mutations along the unique path from the root to the clone in question. For our label-based measurements, we considered the set of mutations first appearing or inherited by each clone. Our two measurements are: (1) the average number of mutations on each node in the entire tree, and (2) a restricted version that computes the average number of such mutations on only the leaves in the tree (therefore implicitly incorporating some topological aspects into the measurement).

#### 3.2.2 Findings from Enumerated Dataset Analysis

Figure 4 shows our results for all 7 of our topological and label-based measurements applied to our 4 datasets of enumerated clonal trees. These results support many trends that we expected to see, and highlight a few that we found surprising.

**Figure 4:**
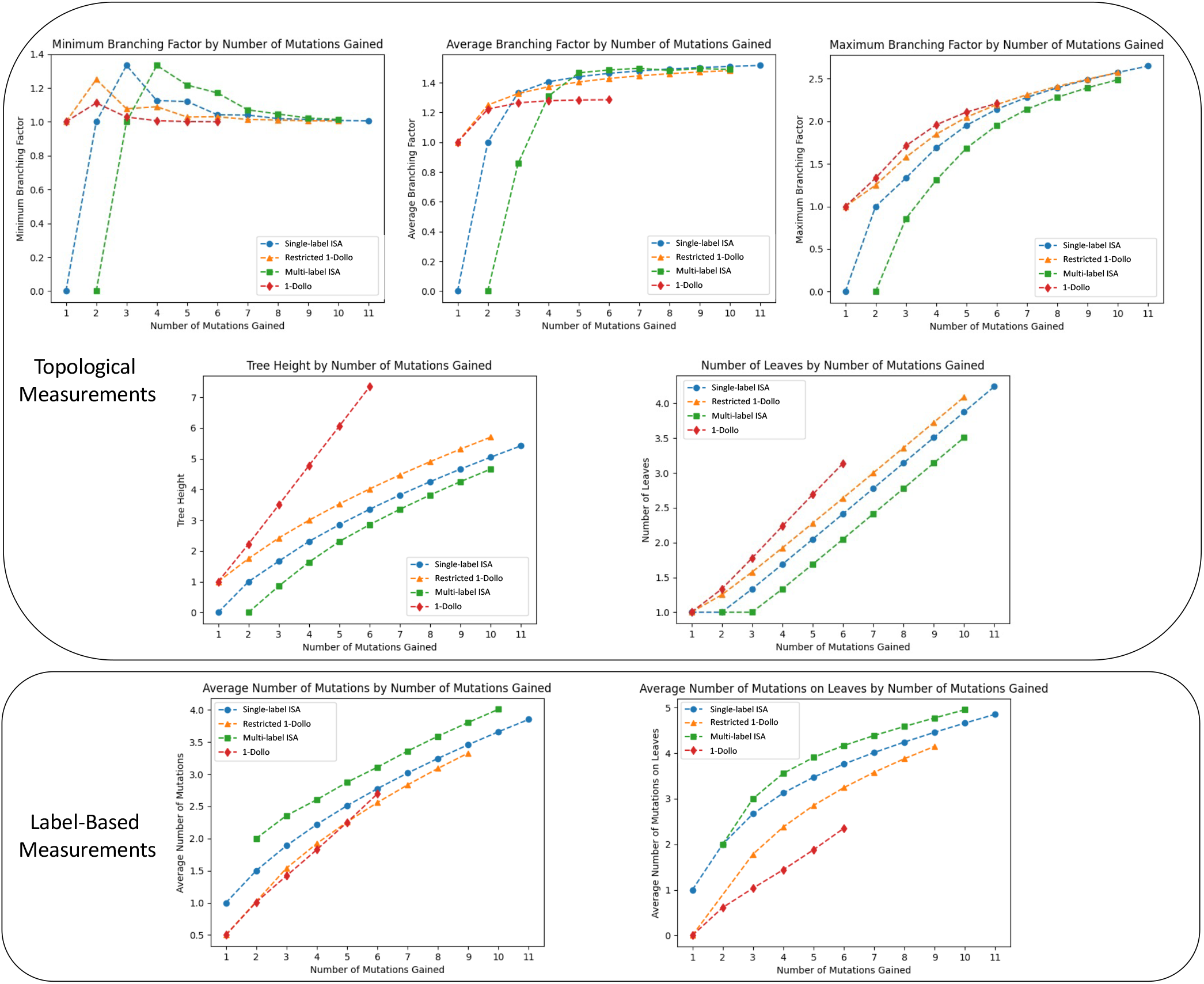
Comparison between four sub-categories of clonal trees. All data points reported are averages across all clonal trees of the specified type with the required number of mutations gained Topological measurements include: minimum, average, and maximum branching factor (top row); tree height and number of leaves (middle row). Label-based measurements include: average number of mutations per node and average number of mutations per leaf (bottom row).

For example, when considering tree height, we see that the single-label ISA trees, multi-label ISA trees, and restricted 1-Dollo trees follow a similar trend as the number of mutations gained increases. The multi-label ISA trees consistently have the smallest height, followed by single-label ISA trees and then the restricted 1-Dollo trees. However, the average height of 1-Dollo trees increases sharply in comparison. This follows from the fact that a 1-Dollo tree with *m* mutations added can also have up to *m* mutations deleted, giving it up to 2*m* nodes, and hence a maximum potential height of 2*m*−1. For instance, when the number of mutations gained is 5 the average height across all 1-Dollo trees is 6.06, much higher than the average height of the single-label ISA trees, whose average height is 2.86. This happens because the 1-Dollo trees can have a height up to 9 (a linear tree where all mutation gains are lost once), compared to single-label ISA trees who have a maximum possible height of 4 (also, a linear tree). This same phenomenon also explains why the restricted 1-Dollo trees have larger heights than the ISA trees (although to a lesser extent).

Another topological measurement that shows an expected pattern is the number of leaves in the trees. We see the same ordering here of the sub-categories of clonal trees as we did with tree height. This time the growth rate for the data we have for all 4 categories appears to grow linearly with the number of mutations gained.

We also see expected trends with our label-based measurements. When considering the average number of mutations on the leaves of the trees, the multi-label ISA trees have the largest values while the 1-Dollo trees have the fewest. All four of the clonal tree categories show similar growth trends as the number of mutations gained increases, although it is difficult to tell whether the 1-Dollo model would continue to follow the same trend for larger numbers of mutations gained. We also see fairly similar trends when considering the number of mutations on just the leaves as when considering all nodes in the tree - just shifted some.

We also see some more unexpected results - in particular with regards to our branching factor analysis. For example, in average branching factor we discovered that single-label ISA trees, multi-label ISA trees, and restricted 1-Dollo trees seemingly converge to a branching factor of 1.6, meaning each non-leaf node has 1.6 children on average. At this time, it is unclear why they converge to this value. 1-Dollo trees have a notably lower average branching factor. We also noticed that minimum branching factor is highest in trees with fewer nodes, rather than those with more, leading to graphs with spikes at the beginning that then fall as more mutations are added. Finally, we were intrigued by the results with regard to maximum branching factor, where all of the metrics seemed to continue rising as more nodes were placed on the tree. On its own this is expected, but what surprised us was that all assumptions seemingly converged to grow together as the number of mutations increased.

### 3.3 Sampled Dataset Creation and Analysis

We created two additional datasets using both the psuedorandom algorithms and Wilson’s algorithm as described in subsection 2.5. These two datasets each contain 1 million trees sampled from the space of single-label ISA trees with 10 mutation gains. We note that the entire space of single-label ISA trees with 10 mutations gained consists of 1 billion trees. So these sampled datasets represent a number of trees that is just 1% of the size of the total space.

We compared these sampled datasets to our enumerated dataset restriction to single-label ISA trees with 10 mutations gains. Specifically, we look at the topological measure of tree height and compared the perfect abundance of trees in each of these three datasets based on the height of trees generated (see Figure 5). Notably, the single-label ISA trees generated using the pseudorandom method were not representative of the actual space of single-label ISA trees. In fact, the pseudorandom method tended to generate much shorter trees (height of 3.62 on average) than what is observed across the entire space of single-label ISA trees (height of 4.84 on average). As expected, Wilson’s algorithm sampled trees whose height distribution is much closer to the actual distribution of all single-label ISA trees (height of 5.05 on average). In fact, the distributions over tree height for Wilson’s algorithm and the entire space of single-label ISA trees look nearly identical, but do indeed have differences. For example, the fraction of trees sampled by Wilson’s algorithm having height 6 is 9.7% whereas the true fraction of all single-label ISA tres with height 6 is 9.68%. This subtle difference is barely perceptible in Figure 5. This experiment makes it clear that Wilson’s algorithm provides samples that are much more reflective of the entire space of single-label ISA trees than does the psuedorandom algorithm.

**Figure 5:**
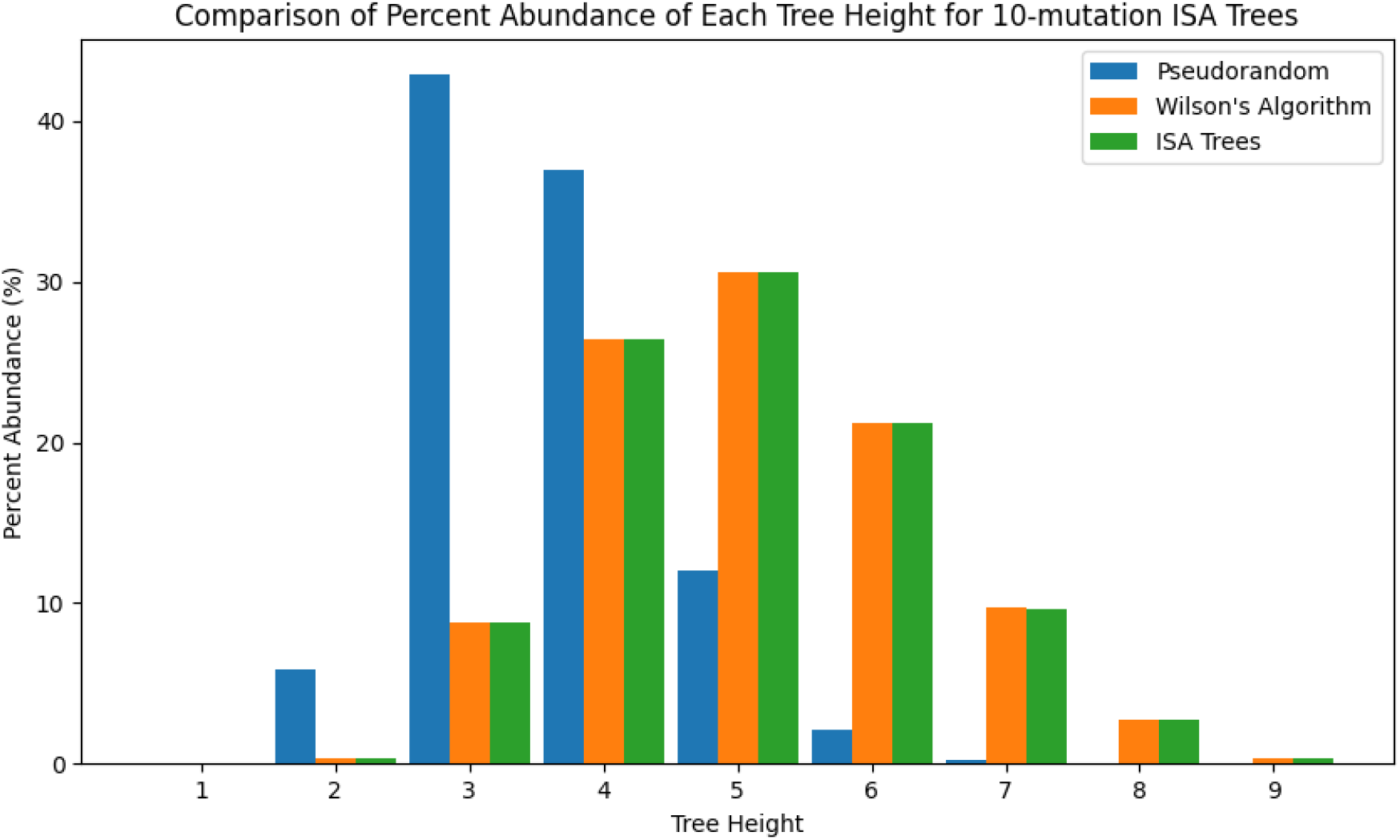
A comparison of the percent abundance of each tree height for 10 mutation single-label IDSA trees sampled using the psuedorandom algorithm and Wilsons algorithm. This is compared the the true percent abundance across the entire enumerated single-label ISA tree dataset. This further serves to demonstrate the uniformity of Wilson’s Algorithm and the lack thereof for the pseudorandom algorithm.

## 4 Conclusion

We have created enumeration algorithms that, for a given number of mutations gained (*m*), generate all single-label ISA, multi-label ISA, restricted 1-Dollo, and 1-Dollo trees. Furthermore, we have shown that it is feasible to generate the entirety of these datasets for small values of *m*. The ability to access the entire clonal tree space for small values of *m* opens up previously unexplored avenues for analyzing and comparing different models of tumor evolution. With modification, our algorithms may be extend to other tumor evolution models, such as the *k*-Dollo model [8] or the Camin-Sokal model [3]. Since the single-label ISA trees contain every topologically distinct tree with *m* labeled nodes, they may be an especially useful starting place for future algorithms to build off.

We used our enumeration algorithms to generate several complete datasets that encompass the entire space of trees under the specified assumption up to a small number of mutations gained. Our analysis of these datasets led to some expected and some unexpected findings and has important implications for how simulated data around these models is created. In particular, new tumor evolution inference methods will want to ensure that any simulated data they are benchmarked on does indeed reflect these entire spaces.

Our findings also enable exploration of some clonal tree spaces for large values of *m*, which may be more biologically relevant than clonal trees with just a few mutations gained. Specifically, our finding that Wilson’s Algorithm returns a representative sample of single-label ISA trees enables the generation and analysis of representative samples of ISA trees with large *m*. This is particularly important when such trees are being used to evaluate the accuracy of new clonal tree inference methods. Of equal importance is our finding that that the method we describe as the pseuodrandom tree sampling method does not provide a representative sample of the space of ISA trees. As a result, this method should be avoided when creating samples of ISA trees that are intended to represent the entire space.

There are many areas for other related future research. In particular, it would be valuable to find or create an algorithm that generates representative samples of restricted 1-Dollo trees and 1-Dollo trees, so that the benefits listed above can apply to other tumor evolution models. In addition, we hope to incorporate recent biological findings about tumor evolution into our measurements and evaluation of the clonal tree space. For instance, researchers have observed that tumor evolution can be differentiated into several different categories, such as linear, branching, punctuated, and neutral [7, 22]. By classifying trees in our four datasets based on their mode of evolution, we could understand how these modes of evolution are represented in different tumor evolution models and whether their representation in the clonal tree space is proportional to the patterns observed in sequencing data. Ultimately, this could help improve tumor evolution models and clonal tree inference methods.

## Acknowledgments

This work has been supported by the National Science Foundation (NSF) award CAREER-IIS-2046011.

## References

[1] Monica-Andreea Baciu-Drăgan and Niko Beerenwinkel. 2024. Oncotree2vec—a method for embedding and clustering of tumor mutation trees. Bioinformatics 40, Supplement_1 (2024), i180–i188.

[2] Leila Baghaarabani, Sama Goliaei, Mohammad-Hadi Foroughmand-Araabi, Seyed Peyman Shariatpanahi, and Bahram Goliaei. 2021. Conifer: clonal tree inference for tumor heterogeneity with single-cell and bulk sequencing data. BMC Bioinformatics 22, 1 (30 Aug 2021), 416. doi:10.1186/s12859-021-04338-7

[3] Joseph H Camin and Robert R Sokal. 1965. A method for deducing branching sequences in phylogeny. Evolution (1965), 311–326.

[4] Arthur Cayley. 1889. A theorem on trees. online. Mathematical papers 13 (1889). http://rcin.org.pl/impan/Content/143722/PDF/WA35_176630_12807-13_Art895.pdf

[5] Simone Ciccolella, Camir Ricketts, Mauricio Soto Gomez, Murray Patterson, Dana Silverbush, Paola Bonizzoni, Iman Hajirasouliha, and Gianluca Della Vedova. 2021. Inferring cancer progression from single-cell sequencing while allowing mutation losses. Bioinformatics 37, 3 (2021), 326–333.

[6] Thomas W Crowther, Henry B Glick, Kristofer R Covey, Charlie Bettigole, Daniel S Maynard, Stephen M Thomas, Jeffrey R Smith, Gregor Hintler, Marlyse C Duguid, Giuseppe Amatulli, et al. 2015. Mapping tree density at a global scale. Nature 525, 7568 (2015), 201–205.

[7] Alexander Davis, Ruli Gao, and Nicholas Navin. 2017. Tumor evolution: Linear, branching, neutral or punctuated? Biochimica et Biophysica Acta (BBA)-Reviews on Cancer 1867, 2 (2017), 151–161.

[8] Louis Dollo. 1893. The laws of evolution. Bull. Soc. Bel. Geol. Paleontol 7 (1893), 164–166.

[9] Mohammed El-Kebir. 2018. SPhyR: tumor phylogeny estimation from single-cell sequencing data under loss and error. Bioinformatics 34, 17 (2018), i671–i679.

[10] Mohammed El-Kebir, Layla Oesper, Hannah Acheson-Field, and Benjamin J. Raphael. 2015. Reconstruction of clonal trees and tumor composition from multi-sample sequencing data. Bioinformatics 31, 12 (06 2015), i62–i70. arXiv:https://academic.oup.com/bioinformatics/articlepdf/31/12/i62/49014148/bioinformatics_31_12_i62.pdf doi:10.1093/bioinformatics/btv261

[11] James S Farris. 1977. Phylogenetic analysis under Dollo’s Law. Systematic Biology 26, 1 (1977), 77–88.

[12] Matthew W Fittall and Peter Van Loo. 2019. Translating insights into tumor evolution to clinical practice: promises and challenges. Genome medicine 11, 1 (2019), 20.

[13] Kiya Govek, Camden Sikes, Yangqiaoyu Zhou, and Layla Oesper. 2022. GraPhyC: Using Consensus to Infer Tumor Evolution. IEEE/ACM Trans Comput Biol Bioinform 19, 1 (Feb. 2022), 465–478.

[14] Ronald L. Graham, Donald E. Knuth, and Oren Patashnik. 1989. Concrete Mathematics: A Foundation for Computer Science. Addison-Wesley, Reading.

[15] Katharina Jahn, Jack Kuipers, and Niko Beerenwinkel. 2016. Tree inference for single-cell data. Genome Biology 17, 1 (05 May 2016), 86. doi:10.1186/s13059-016-0936-x

[16] Motoo Kimura. 1969. The number of heterozygous nucleotide sites maintained in a finite population due to steady flux of mutations. Genetics 61, 4 (1969), 893.

[17] Donald E Knuth. 2011. The art of computer programming, volume 4A: combinatorial algorithms, part 1. Pearson Education India.

[18] Jack Kuipers, Katharina Jahn, Benjamin J Raphael, and Niko Beerenwinkel. 2017. Single-cell sequencing data reveal widespread recurrence and loss of mutational hits in the life histories of tumors. Genome research 27, 11 (2017), 1885–1894.

[19] Jiaying Lai, Yi Yang, Yunzhou Liu, Robert B Scharpf, and Rachel Karchin. 2024. Assessing the merits: an opinion on the effectiveness of simulation techniques in tumor subclonal reconstruction. Bioinformatics Advances 4, 1 (06 2024), vbae094. https://academic.oup.com/bioinformaticsadvances/article-pdf/4/1/vbae094/58362429/vbae094.pdf doi:10.1093/bioadv/vbae094

[20] Lin Li, Wenqin Xie, Li Zhan, Shaodi Wen, Xiao Luo, Shuangbin Xu, Yantong Cai, Wenli Tang, Qianwen Wang, Ming Li, et al. 2024. Resolving tumor evolution: a phylogenetic approach. Journal of the National Cancer Center (2024).

[21] Magda Markowska et al. 2022. CONET: copy number event tree model of evolutionary tumor history for single-cell data. Genome Biology 23, 1 (2022), 128.

[22] Robert Noble, Dominik Burri, Cécile Le Sueur, Jeanne Lemant, Yannick Viossat, Jakob Nikolas Kather, and Niko Beerenwinkel. 2022. Spatial structure governs the mode of tumour evolution. Nature ecology & evolution 6, 2 (2022), 207–217.

[23] Peter C Nowell. 1976. The Clonal Evolution of Tumor Cell Populations: Acquired genetic lability permits stepwise selection of variant sublines and underlies tumor progression. Science 194, 4260 (1976), 23–28.

[24] Heinz Prüfer. 1918. Neuer beweis eines satzes über permutationen. Arch. Math. Phys 27, 1918 (1918), 742–744.

[25] Edith M. Ross and Florian Markowetz. 2016. OncoNEM: inferring tumor evolution from single-cell sequencing data. Genome Biology 17, 1 (15 Apr 2016), 69. doi:10.1186/s13059-016-0929-9

[26] Sohrab Salehi, Fatemeh Dorri, Kevin Chern, Farhia Kabeer, Nicole Rusk, Tyler Funnell, Marc J. Williams, Daniel Lai, Mirela Andronescu, Kieran R. Campbell, Andrew McPherson, Samuel Aparicio, Andrew Roth, Sohrab P. Shah, and Alexandre Bouchard-Côté. 2023. Cancer phylogenetic tree inference at scale from 1000s of single cell genomes. Peer Community Journal 3, Article e63 (2023). doi:10.24072/pcjournal.292

[27] Palash Sashittal, Haochen Zhang, Christine A Iacobuzio-Donahue, and Benjamin J Raphael. 2023. ConDoR: tumor phylogeny inference with a copy-number constrained mutation loss model. Genome biology 24, 1 (2023), 272.

[28] N.J.A. Sloane. 2014. A Handbook of Integer Sequences. Elsevier Science. https://books.google.com/books?id=pw3jBQAAQBAJ

[29] Maitena Tellaetxe-Abete, Charles Lawrie, and Borja Calvo. 2025. Addressing the multiplicity of optimal solutions to the Clonal Deconvolution and Evolution Problem. European Journal of Operational Research 320, 3 (2025), 777–788. doi:10.1016/j.ejor.2024.09.006

[30] Xiaodong Wang, Lei Wang, and Yingjie Wu. 2009. An Optimal Algorithm for Prufer Codes. J. Softw. Eng. Appl. 02, 02 (2009), 111–115.

[31] David Bruce Wilson. 1996. Generating random spanning trees more quickly than the cover time. In Proceedings of the Twenty-Eighth Annual ACM Symposium on Theory of Computing (Philadelphia, Pennsylvania, USA) (STOC’96). Association for Computing Machinery, New York, NY, USA, 296–303. doi:10.1145/237814.237880

[32] Hamim Zafar, Nicholas Navin, Ken Chen, and Luay Nakhleh. 2019. SiCloneFit: Bayesian inference of population structure, genotype, and phylogeny of tumor clones from single-cell genome sequencing data. Genome research 29, 11 (2019), 1847–1859.

[33] Hamim Zafar, Anthony Tzen, Nicholas Navin, Ken Chen, and Luay Nakhleh. 2017. SiFit: inferring tumor trees from single-cell sequencing data under finite-sites models. Genome biology 18 (2017), 1–20.

